# XTree enables memory-efficient, accurate short- and long-read pseudoalignment to large genome databases

**DOI:** 10.64898/2025.12.22.696015

**Authors:** Gabriel A Al-Ghalith, Victor Boddy, Krista A Ryon, James R Henriksen, David C Danko, Brett Farthing, Matteo Marengo, George M Church, Chirag J Patel, Dan Knights, Braden T Tierney

## Abstract

XTree is a k-mer-based aligner enabling rapid, memory-efficient alignment of sequencing reads to large whole-genome reference databases. Here, we detail XTree’s performance on short- and long-read sequencing data and demonstrate its high accuracy across diverse bacterial and archaeal genomes. In benchmarks against five widely used taxonomic profilers on synthetic prokaryotic metagenomes built from genomes absent from the reference database, XTree achieved the highest species-level F1 score for both short reads (0.899) and long reads (0.938), while classifying reads at approximately 2 s per million short reads and 30 s per million long reads, 4- to 300-fold faster than the other tools. We additionally provide a pre-indexed database for the entire Genome Taxonomy Database (version r232, compression level 4).

## Main Text

The alignment of next-generation sequencing (NGS) data to reference genomes is foundational to modern genomics. Mapping bulk sequencing reads to large databases of known genomes underpins our ability to quantify the genomic and taxonomic diversity of complex ecosystems. However, alignment and classification algorithms designed many years ago do not necessarily (1) computationally scale to today’s large reference databases; (2) guarantee high sensitivity and specificity with continually increasing species diversity across the domains of life; or (3) perform well on newer sequencing methods (e.g., long-read sequencing).

Here, we present XTree and its accompanying databases. XTree (see *Methods*) is a k-mer-based pseudoaligner designed for memory-efficient (including out-of-core) parallel pseudoalignment and classification of long or short sequencing reads against large reference genome databases. Although XTree’s predecessors (e.g., BURST, UTree, XTree V1.0) have been used in both academia and industry over the past decade^1–10^, the XTree algorithm itself has not previously been described or benchmarked against comparison tools in a peer-reviewed manuscript. Accordingly, in this report, we formally detail a new version of XTree (V2.2.13) for the first time in the scientific literature.

XTree builds on prior k-mer-based genome indexing and read assignment strategies^11,12^ (Fig 1A) by using: (1) memory mapping for efficient database usage by simultaneous processes; (2) motif- and hash-based k-mer subsampling (as opposed to using minimizers^13^); (3) canonicalization of reference k-mers into a binary search tree for fast lookups; and (4) optional winner-take-all read redistribution, which, for each read, takes the references tied for the highest matched k-mer count and resolves the tie using the support for those references across the whole sample.

**Figure 1:**
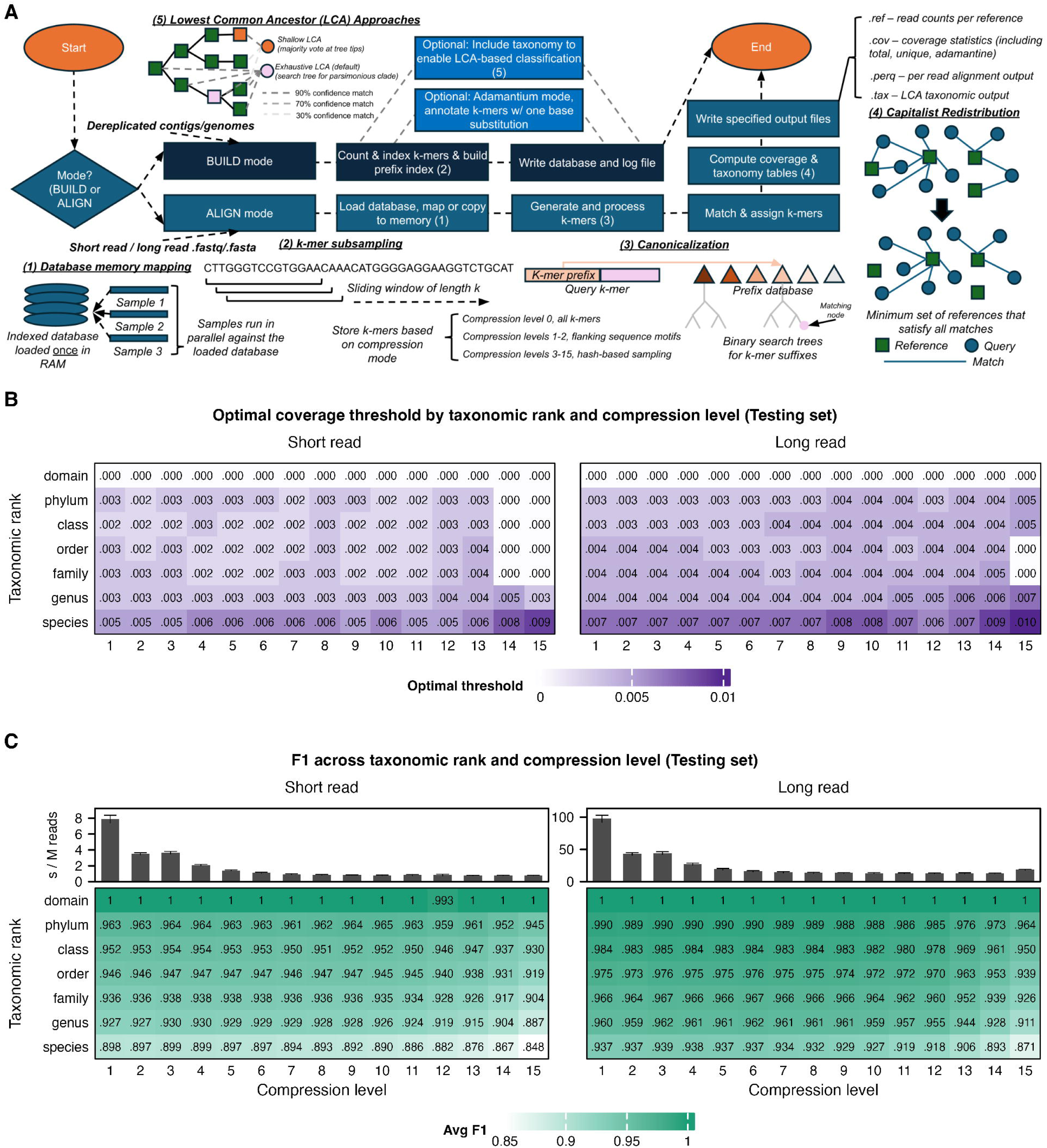
XTree overview and performance on short- and long-read synthetic prokaryotic metagenomes (testing set). A) Overview of the XTree algorithm. XTree utilizes k-mer-based indexing with motif- or hash-based subsampling, canonicalization via binary search trees, and optional “capitalist” read redistribution for read assignment, as well as multiple options for Lowest Common Ancestor (LCA) analysis. This figure diagrams motif-based sampling at compression levels 1–2. Levels 3–15 sample canonical k-mers by their SplitMix64 hash. Level 0 keeps all valid k-mers. B) Coverage threshold that maximizes mean F1 score at each taxonomic rank and compression level for short and long reads. C) Mean F1 score by taxonomic rank and compression level for the 50 testing metagenomes (non-representative genomes absent from the database) at a 1% coverage threshold, for short and long reads; bars above each heatmap show the mean (± SD) alignment time per million reads.

XTree additionally identifies multiple types of k-mers and reports associated coverage “types” with each: unique k-mers are those found in only a single genome, whereas Adamantine (“highly unique”) k-mers are those that are a Hamming distance of > 1 (i.e., more than one base substitution) away from all k-mers in other references, enabling strain-specific classification. By default, XTree reports total coverage of a genome (proportion of an entire genome covered), unique coverage (unique regions of a genome covered), and Adamantine coverage (Adamantine regions of a genome covered).

To showcase XTree’s utility and guide its use, we evaluated its performance across various synthetic metagenomes. Beginning with prokaryotes, we randomly selected 20,000 Genome Taxonomy Database (GTDB)^14^ species clusters, each with a representative genome and at least one non-representative genome, and indexed the representative genomes. This allowed us to create both a training set of 50 synthetic metagenomes composed of the same genomes used to build the database, and a testing set of 50 synthetic metagenomes composed of non-representative genomes from the same species clusters, i.e., new genomes not present during database indexing. Because the testing set better reflects real samples, in which most strains are absent from reference databases, we present testing-set results in the main figures and training-set results in the Supplement. In a sweep of k-mer lengths (k = 21, 24, and 29, each at compression levels 2, 4, 8, and 12), performance was similar across k-mer lengths (Supp Fig 1). Using k=29 yielded the highest mean F1 score at the maximized coverage threshold (0.953 versus 0.951 for k = 24 and 0.940 for k=21; short reads, average over compression levels and both training and testing sets), although at the fixed 1% threshold, k =24 and k=29 were indistinguishable.

All subsequent analyses used XTree v2.2.13 with k = 29, evaluated at each compression level. Next, we evaluated performance across taxonomic ranks to determine the effects of compression level and coverage threshold on mean F1 scores. On the testing set, the highest species-level mean F1 score was 0.920 at a coverage threshold of 0.5% for short reads and 0.950 at a threshold of 0.7% for long reads (both at compression level 3), and the F1-maximizing threshold was at or below 1% at every taxonomic rank and compression level (Fig 1B; training set in Supp Fig 2B). We therefore used a fixed 1% threshold, which performed well across all compression levels, for the rest of the analysis. At this threshold, the species-level mean F1 score on the short-read testing set remained between 0.890 and 0.899 from compression level 1 to 10 (mean precision 0.926–0.940; mean recall 0.857–0.866) before declining to 0.848 at compression level 15, and was higher at broader ranks (e.g., 0.930 at the genus and 0.964 at the phylum level at compression level 4; Fig 1C). The training set followed the same pattern at higher values, with a species-level mean F1 score of 0.968–0.975 (precision 1.000) up to compression level 10, declining to 0.928 at compression level 15 (Supp Fig 2A). These results showcase XTree’s ability to accurately assign species-level classifications to genomes not included during database construction. Meanwhile, higher compression levels substantially reduced resource use: from compression level 1 to 10, database size fell from 156 GB to 1.5 GB, and mean short-read alignment time fell from 7.9 s to 0.8 s per million reads (Fig 1C).

**Figure 2:**
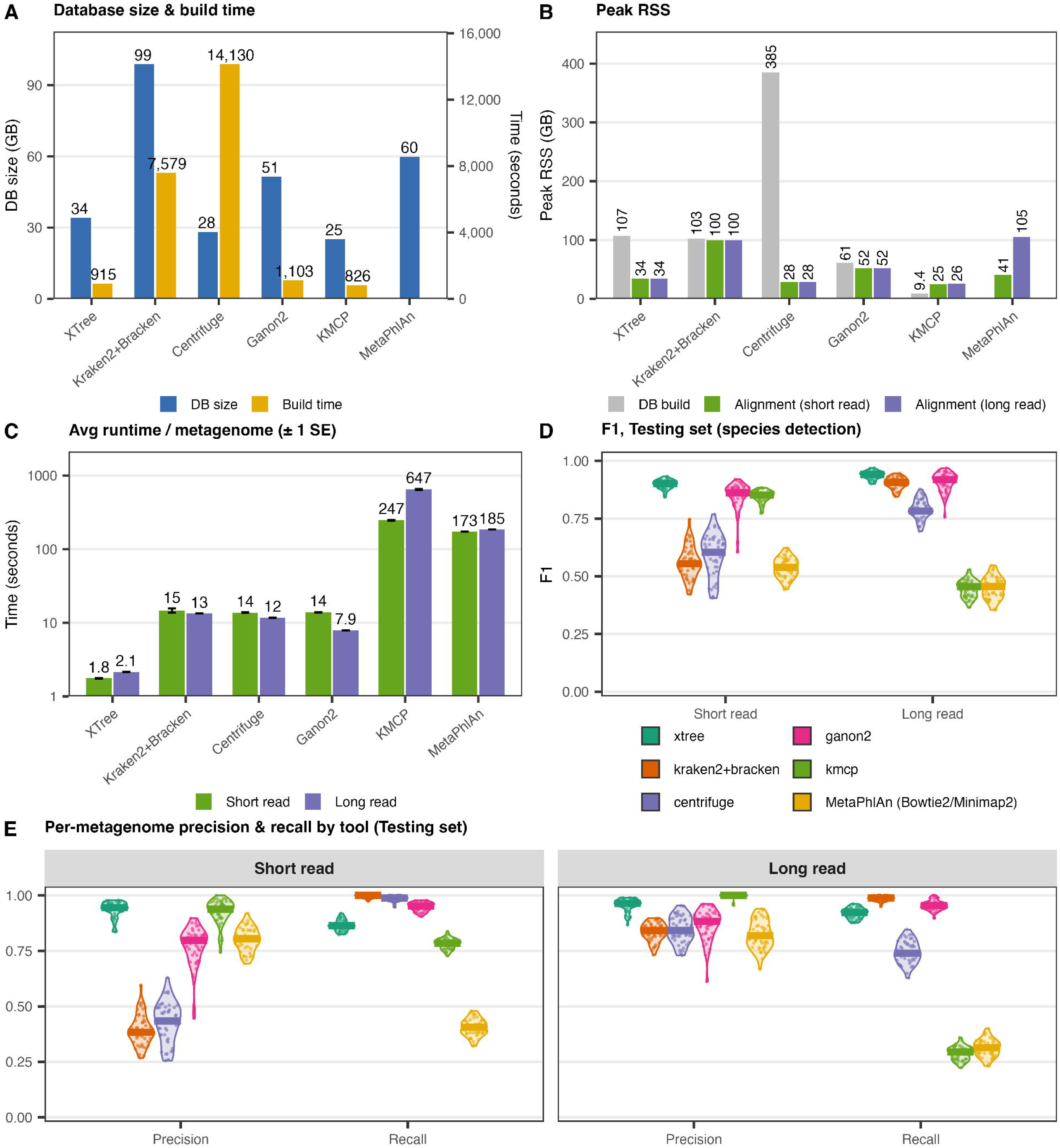
Comparison of XTree with other metagenomic profilers on synthetic prokaryotic metagenomes. A) Database size and build time. B) Peak memory (resident set size) during database build and alignment. C) Mean alignment or classification time per metagenome (± SE) for short- and long-read metagenomes. D) Per-metagenome species-level F1 score on the testing set for short- and long-read metagenomes, horizontal line marks the median. E) Per-metagenome precision and recall on the testing set for short and long reads; each point is one metagenome and horizontal lines mark the median. XTree results use the compression level 4 database (k = 29) and a 1% coverage threshold; the other tools were run with default settings. MetaPhlAn used its default marker database, with Bowtie2 for short reads and Minimap2 for long reads; KMCP runtime is the sum of its search and profile steps. Training-set results are shown in Supp Fig 2C–D.

With accurate prokaryotic species detection established, we next evaluated XTree’s relative abundance estimates by comparing observed and expected relative abundances at each compression level. On the testing set, the Pearson correlation between expected and observed relative abundance was 0.930–0.934 for short reads and 0.930–0.933 for long reads up to compression level 13, falling only to 0.922 and 0.926, respectively, at compression level 15 (Supp Fig 3A). Agreement was near-perfect on the training set (Pearson r ≥ 0.999 for short reads and ≥ 0.997 for long reads up to compression level 12, and 0.993 and 0.991, respectively, at compression level 15; Supp Fig 3B). The weaker correlation on the testing set likely reflects that non-representative genomes share only part of their sequence with the representative genomes in the database.

**Figure 3:**
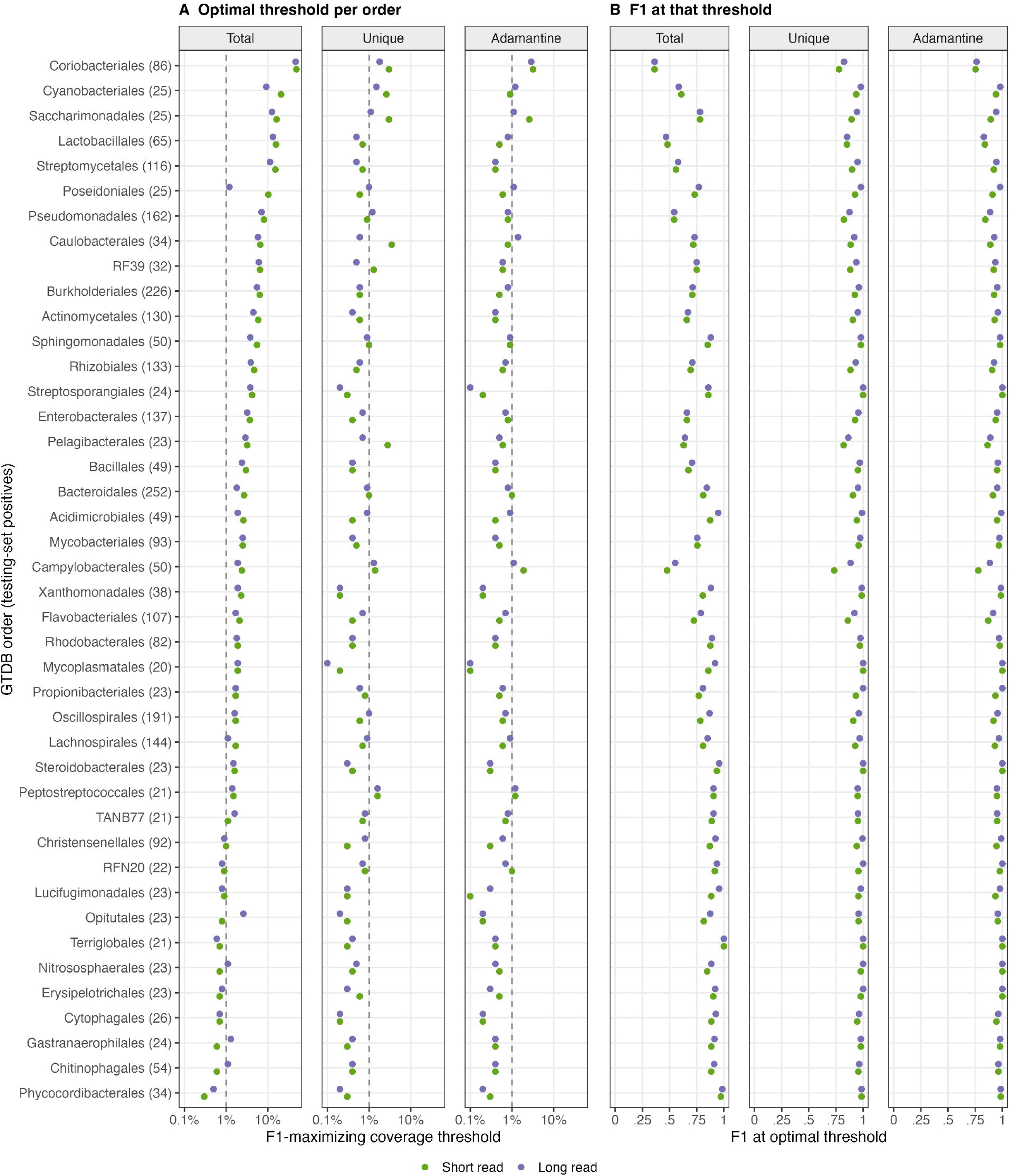
Per-order coverage thresholds. A) Coverage threshold that maximizes F1 for each GTDB order, with total, unique, and Adamantine coverage each used alone to call a genome present (log scale). B) F1 score at that threshold. Results are for short reads (green) and long reads (purple) for the testing set at compression level 4 (XTree v2.2.13, k=29). The 42 orders shown are those with at least 20 positive genome-metagenome pairs; that number is given in parentheses. Within each order, true and false positives were pooled across the 50 testing metagenomes. Thresholds were swept from 0 to 100% in 0.1% steps, and the lowest threshold that maximized F1 score was taken. The dashed line marks the 1% threshold used throughout. Under the detection rule used in all benchmarks (min(total coverage, unique coverage, Adamantine coverage)), per-order optima had a median of 0.5%.

We next compared XTree with other commonly used metagenomic taxonomic profilers. We selected Kraken2+Bracken, Ganon2, Centrifuge, KMCP, and MetaPhlAn4 (which uses Bowtie2 to align short reads) to compare both classification performance and hardware requirements. Similar to XTree, Kraken2, Ganon2 and KMCP each employ k-mer-based strategies^11,17,18^, whereas Centrifuge uses an FM-index-based approach^19^ and MetaPhlAn4 aligns reads to clade-specific marker genes^20^. The indexing and classification algorithms of each tool are summarized in Table 1. Kraken2, Ganon2, Centrifuge, and KMCP indexed the same representative genomes as XTree, MetaPhlAn used its default marker database, and each tool profiled both sets of 50 training and testing metagenomes; for XTree, we used the compression level 4 database (the v2.2.13 default) and the 1% coverage threshold.

**Table 1:** Indexing and classification approaches of the six tools compared on the synthetic prokaryotic benchmark (Fig 2; Supp Fig 2). Except for MetaPhlAn, which uses its own marker-gene database, every tool built its index from the same 19,999 GTDB Release 232 representative genomes. Versions are those used in this study.

| Tool | Version | Index / database | Classification approach |
|---|---|---|---|
| XTree | v2.2.13 | Prefix-hashed, suffix-sorted k-mer index (binary search) of whole genomes (k = 29; compression level 4); memory-mapped and shared between processes | Exact k-mer matching of reads to reference genomes; each read assigned to its best-supported reference; genome presence called from total, unique and Adamantine breadth of coverage (1% threshold) |
| Kraken2 + Bracken | Kraken2 2.17.1; Bracken 3.1 | Compact hash table of minimizers (k = 35, minimizer length 31) with GTDB taxonomy; Bracken read-length distributions for 150 bp (short) and 4,000 bp (long) reads | Minimizer hits mapped to taxa and each read assigned to the highest-scoring root-to-leaf path (lowest common ancestor); Bracken re-estimates species abundance with a Bayesian model |
| Centrifuge | 1.0.4.2 | FM-index (Burrows–Wheeler transform) of the reference genomes | Variable-length exact matches between reads and the index are scored; each read assigned to its best-scoring taxa, collapsed to their lowest common ancestor when there are too many |
| Ganon2 | 2.3.0 (short reads); 2.4.1 (long reads) | Hierarchical interleaved Bloom filter (HIBF) of k-mers from the reference genomes | Reads matched by shared k-mer counts against the filter; multi-matching reads reassigned with an expectation-maximization algorithm to estimate abundance |
| KMCP | 0.9.5 | Modified COBS (compact bit-sliced signature) index of k-mers from genome chunks | Reads searched by k-mer containment against genome chunks; profiling filters candidate genomes by chunk coverage and estimates abundance |
| MetaPhlAn | 4.2.5 (database mpa_vJan26_CHOCOPhlAnSGB_202605) | Stock clade-specific marker-gene database (not built from the benchmark genomes) | Reads aligned to marker genes (Bowtie2 2.5.5 for short reads; Minimap2 for long reads); species abundance estimated from marker coverage and mapped to GTDB for scoring |

On the testing set, XTree achieved the highest mean species-level F1 score (0.899), followed by Ganon2 and KMCP (both 0.848), Centrifuge (0.590), Kraken2+Bracken (0.560), and Bowtie2+MetaPhlAn (0.537) (Fig 2D). Kraken2+Bracken and Centrifuge detected nearly every species in the testing set but with low precision (median 0.384 and 0.434, respectively), whereas XTree maintained a median precision of 0.945 with a median recall of 0.864 (Fig 2E). On the training set, in which every genome is present in the database, Kraken2+Bracken and Ganon2 achieved slightly higher F1 scores than XTree (0.993 versus 0.974), as XTree’s perfect precision was offset by a lower recall (0.950; Supp Fig 2C–D). Next, XTree’s database (34 GB) was similar in size to those of Centrifuge (28 GB) and KMCP (25 GB) and smaller than those of Ganon2 (51 GB), MetaPhlAn (60 GB), and Kraken2 (99 GB), and XTree and KMCP had the fastest build times (915 s and 826 s, respectively, versus 1,103–14,130 s for the other tools; Fig 2A). Although database construction occurs only once, XTree’s advantage was most apparent in average alignment time per metagenome: 1.8 s, versus 14–15 s for Kraken2+Bracken, Centrifuge, and Ganon2, and 173 s and 247 s for Bowtie2+MetaPhlAn and KMCP, respectively (Fig 2C). XTree’s peak memory during alignment (34 GB, nearly all of which was the shared, memory-mapped database) was lower than that of Kraken2+Bracken (100 GB) and Ganon2 (52 GB) but higher than that of Centrifuge (28 GB) and KMCP (25 GB), and its database build peaked at 107 GB of RAM, compared with 9 GB for KMCP and 385 GB for Centrifuge (Fig 2B).

We next characterized XTree’s performance on long-read data. As long-read sequencing becomes more affordable and widespread, new bioinformatic tools should perform well on long reads. As with short reads, performance in the k-mer sweep was similar for k = 21–29, with k = 29 performing best (Supp Fig 1). Mean F1 scores were higher than for short reads at all compression levels: on the testing set, the species-level mean F1 score was 0.927–0.939 up to compression level 10 (mean precision 0.941–0.959; mean recall 0.914–0.921), declining to 0.871 at compression level 15 (Fig 1C), and on the training set it was 0.981–0.986 up to compression level 10 (Supp Fig 2A). Mean long-read alignment time decreased from 97 s to 13 s per million reads from compression level 1 to 10 (Fig 1C). Next, we compared the long-read performance of XTree to the same suite of tools, using Minimap2^21^ in place of Bowtie2 for MetaPhlAn, as Bowtie2 does not support long-read data. On the testing set, XTree again achieved the highest mean F1 score (0.938), followed by Ganon2 (0.908), Kraken2+Bracken (0.903), Centrifuge (0.790), Minimap2+MetaPhlAn (0.450), and KMCP (0.447) (Fig 2D). As with short reads, several tools slightly exceeded XTree on the training set (Centrifuge, 0.998; Ganon2, 0.997; Kraken2+Bracken, 0.994; XTree, 0.986; Supp Fig 2C–D). XTree classified each long-read metagenome in 2.1 s on average, 3.7- to 300-fold faster than the other tools (7.9 s for Ganon2 to 647 s for KMCP; Fig 2C).

We also performed a brief analysis of archaeal genomes. GTDB Release 232 contained 3,013 archaeal species clusters with at least one representative and one non-representative genome; as for the prokaryotic benchmark, we indexed the representative genomes (3,012 in the final database, because one genome failed to download) with XTree v2.2.13 at k = 29 at each of compression levels 1–15, and generated 50 training and 50 testing synthetic metagenomes with both short- and long-read data (see Methods). On the testing set at a 1% coverage threshold, the species-level mean F1 score was 0.884–0.896 for short reads and 0.928–0.940 for long reads up to compression level 10, declining to 0.834 and 0.850, respectively, at compression level 15; on the training set it was 0.961–0.970 for short reads and 0.986–0.992 for long reads up to compression level 10 (Supp Fig 4A–B). The F1-maximizing coverage threshold was at or below 1.2% at every taxonomic rank and compression level (Supp Fig 4C–D); averaged over compression levels and both sets, the thresholds that maximized mean F1 score were 0.6% for short reads and 0.8% for long reads. Again, we used 1% because it performed consistently well across all compression levels, and we recommend it for taxonomic classification of both bacterial and archaeal samples. Because the archaeal reference set is much smaller, the compression level 4 database was 3.9 GB and short-read alignment took 1.8 s per million reads on average. We next asked whether a single detection threshold generalizes across prokaryotic clades. For each GTDB order with ≥20 true-positive genomes, we identified the F1-optimal threshold on the minimum of XTree’s three reported proportions. Optimal thresholds were consistently low (median 0.5%, range 0.1–3.0%; all orders <10%), and closely tracked those of unique coverage alone, which is typically the limiting proportion (Fig 3A). In contrast, total coverage alone yielded widely varying order-specific optima (0.3–49%) and poor discrimination (pooled F1 0.48), reflecting k-mer sharing between related genomes. Although per-order optima were typically below 1%, we used a single 1% threshold as a conservative default across all orders. Compared with each order’s optimal threshold, it reduced false positives by 22% and 26% for short reads and long reads, at a cost of a median F1 score decrease of 0.03 and 0.02, respectively. Thus, a single low threshold on XTree’s combined coverage metric supports presence/absence calls across diverse prokaryotic clades without clade-specific tuning.

In summary, we describe XTree, an aligner built for robust and flexible whole-genome-based classification across diverse use-cases. The codebase, as well as download links for all databases and synthetic data in this study, are fully available at https://github.com/two-frontiers-project/2FP-XTree. We anticipate that these tools will be broadly useful to the scientific community.

## Supplemental

## Methods

### XTree databases and indexing

XTree is a k-mer-based aligner. It matches reference sub-sequences of length k – determined during the database build step – to k-mers found in input sequences. Input sequences can be in FASTQ or FASTA format, paired- or single-end, and of any length. The general process for the XTree algorithm, both database construction and alignment, is described in Fig. 1A.

Databases can be constructed of any set of contigs in FASTA format. We recommend high-quality genomes that are dereplicated at a level between 90% and 99% nucleotide identity (e.g., 95% for bacterial genomes). We do not recommend constructing databases from multiple domains of life, as the optimal k-mer length or dereplication cutoff may vary depending on genome size and complexity. Genome dereplication is important prior to database generation as it enables the identification of species- or strain-specific k-mers, which in turn informs a user’s ability to design coverage cutoffs.

Databases are built by subsampling k-mers according to a user-selected compression level (0–15; default 4 in v2.2.13), which determines the number of k-mers retained in the index. For compression level 0, all k-mers of a given length are stored, resulting in the largest databases. At compression levels 1 and 2, sequence motifs determine which k-mers are preserved in the index (sampling stride); the motifs in question were chosen to capture a wide range of genomic sequences while reducing the redundancy of stored information (i.e., minimizing tightly overlapping k-mers while retaining comparability of the same k-mers regardless of context). At compression levels 3–15, k-mers are instead retained by hash-based subsampling (splitmix64 hash^22^), with each successive level retaining approximately half as many k-mers as the previous level; hash-based subsampling can optionally be applied to levels 1 and 2 as well (--enable-hashbrowns), which was not used in this study. For example, the k = 29 prokaryotic benchmark database stored 19.5 billion k-mers (156 GB) at compression level 1, 4.1 billion (34 GB) at the default compression level 4, and 2.0 million (1.0 GB) at compression level 15. During alignment, query k-mers are subsampled in the same way as the database.

XTree currently stores all k-mers that meet these subsampling criteria, and it additionally can track k-mers that appear in a single reference as being “unique” and/or “Adamantine.” Unique k-mers are those found in a single genome. Adamantine k-mers are a subset of these; each Adamantine k-mer differs by more than one base substitution (Hamming distance of 2 or greater) from all k-mers present in other genomes; this makes Adamantine k-mers a more reliable indicator of that specific reference, rather than a simple off-by-one sequencing error/small mutation/noise flip. We propose that they are useful for tracking strains when multiple similar strains are present, as sequencing noise can often flip a k-mer to appear like it’s from another similar genome. We do not recommend using Adamantine k-mers in cases where strain-level specificity is not necessary.

During alignment, query k-mers are canonicalized into a binary format consisting of a binary prefix (for use as initial perfect hash index) and a suffix, which is looked up within a binary tree stored at that prefix. We use a flattened binary tree due to its efficiency of data structure, with lower RAM usage than a full hash map given its ability to assemble full k-mers on the fly using only their stored suffixes combined with the tree structure. Effectively, lookup consists of an initial perfect hash lookup of the binary prefix, which provides the index range within the flattened tree array containing suffix-keyed leaves, within which the (first) suffix match is identified by left-first binary search (LBS) within that range.

The completed lookup provides an index into all matching genomes and associated metadata (e.g., source genome identifier, the source genome’s taxonomy, etc). If present in the database structure, a secondary “Adamantine” tree is also searched by default for each unique k-mer to determine whether the k-mer is “weakly unique” (i.e., found within a Hamming distance of 1 of a k-mer in another genome); if a unique k-mer is not specified as weakly unique, it is Adamantine (strongly unique). In v2.2.13, this weak-uniqueness test is applied by default against every reference k-mer on both strands, including k-mers not retained by subsampling; the faster heuristic test from earlier versions (v2.2.6), which probes only the one-substitution neighbours of stored k-mers, remains available at build time (--fast-adamantium). All databases benchmarked here used the default (full) test. The Adamantine tree is generated by default on database construction, though, as mentioned above, for analyses not reliant on strain-level differentiation, it can be omitted during the database build step (or not utilized during alignment even if present in the database), enabling downstream efficiency gains during alignment.

If a taxonomy map is provided in the database BUILD process (or at alignment time with --map), two forms of taxonomy output are available in ALIGN mode using the --tax-out flag. Using the flag will default to using Lowest Common Ancestor (LCA) taxonomic assignment by performing a root-to-leaf interpolation among each query’s set of identified matches, adhering to the user-specified confidence (--confidence) level at each level of the hierarchy; a simple leaf-only voting scheme will be used instead if --shallow-lca is specified^23^. Alternatively, the Capitalist read redistribution algorithm will be used for both taxonomy assignment and reference re-assignment if --redistribute is specified (up to seven refinement passes by default, adjustable with -- redistribute-passes), similar to its use in BURST but using an expectation maximization step to approximate BURST’s graph-pruning method for simplicity^1^. When Capitalist is enabled with -- tax-out, taxonomy will be reported at the leaf node of the winning taxonomic label (highest specificity).

During the alignment process, a given database is mapped to memory, enabling multiple XTree processes on the same machine to simultaneously use a single copy of the loaded database in dynamic shared memory backed by a single location on physical disk, reducing overall memory footprint, enabling “warm starts” for subsequent runs, and allowing for simultaneous instances to run. Small databases can alternatively be copied to CPU-local memory (--copymem), providing each process greater locality of reference; on some architectures (such as multi-socket systems for which NUMA controls are not enabled) or for some very small databases that can partially fit within L3 cache, this may be preferred. We note that as CPU-local memory sizes increase, XTree’s performance will increase for proportionally larger databases. Notably, due to memory mapping, XTree does not strictly require the entire database to fit within RAM at one time. Users with limited memory (as low as <= 64GB) but with fast direct NVME SSD storage may run XTree without issue but incur a speed penalty approximately proportional to both the difference between the size of the database and the free RAM, and the speed differential between the SSD random read performance and system RAM bandwidth.

### XTree alignment, read assignment, and memory efficiency

Upon ingestion of reads, XTree begins its k-mer search in the same manner as it approaches database indexing. All k-mers for a given read that pass the same subsampling as the reference database (motif- or hash-based, depending on its compression level) are stored in memory. These k-mers are matched to the canonicalized reference database and the appropriate leaves of the binary search tree are searched. Each potential match to a given reference genome is stored.

When --redistribute is specified, XTree runs in “Capitalist Redistribution” (i.e., winner takes all) mode, where the references tied for the highest number of matched k-mers are retained for each read, and each tie is resolved using the candidates’ support across the entire sample. Capitalist is a global read redistribution algorithm, as opposed to a per-read greedy assignment method. Its purpose is to resolve multi-mapping reads across the entire dataset simultaneously, as opposed to resolving each query independently. It leverages the fact that XTree records all possible reference hits for every k-mer in every read, producing a large, multi-mapped space describing which reads could feasibly derive from which sources. Capitalist performs an expectation-maximization procedure over this space; in each iteration, each read is provisionally assigned to whichever of its tied top references is most strongly supported by other reads across the entire dataset. As assignments aggregate, the relative support for competing references shifts accordingly. This process continues until convergence on a given reference, or until an iteration limit is reached (seven passes by default), producing a one-to-one assignment in which a read is assigned to a single reference. The phrase “winner takes all” refers to this globally considered outcome, as opposed to a per-read or per-reference approach. Therefore, capitalist redistribution is approximating the solution to a global set-cover-like problem without explicitly solving an NP-complete formulation, using dataset-wide competition among references to converge on a parsimonious explanation across all reads, jointly.

This behavior is distinct from other approaches (e.g., fast read mappers or Kraken2), which rely on per-read best-hit (or stored LCA-based) assignment modes. Other read redistribution techniques (e.g. Bracken) use post hoc abundance redistribution without access to information on each read’s specific multi-mappings, and do not explicitly leverage the set of multi-mappings across all queries in a dataset for making assignment decisions. If Capitalist Redistribution mode is disabled (the default), the reference genome with the highest number of k-mer matches is simply assigned (with “first stored reference” as a tie breaker).

If XTree is run with the --tax-out command (without Capitalist Redistribution), XTree will independently compute the LCA among all reference genomes matched by the query. Importantly, LCA is not stored per reference k-mer based on a pre-computed database; it is computed among the specific matches for each query during alignment. Specifically, the LCA taxonomic assignment of a read is computed after the k-mer lookup stage by traversing the taxonomic subtree formed by all references with matching k-mers in the read; if --tax-out is run with Capitalist Redistribution, the taxonomic call of a read is also determined based on the full-dataset-aware read assignment on the taxonomic string provided; in tandem with the reference-level capitalist redistribution, this enables multiple independent granularities of assignment for a single query, leveraging the different scales of evidence available at different set granularities.

### XTree outputs and coverage thresholding

In addition to the timing and alignment statistics printed to stdout by default, XTree generates up to five output files depending on user specification: tax, cov, ref, perq, and orthog. The tax file, as previously described, contains taxonomic information in a simple tab-delimited format of key-value association (reference, followed by semicolon-delimited hierarchical string). To use “--tax-out” mode, a taxonomy map must be supplied either during database construction or at alignment time (--map, which overrides any map embedded in the database). The taxonomy file may contain up to two hierarchies (e.g., two different taxonomic schemas or granularities, or a taxonomy and a GO hierarchy if genes are used as references instead of genomes), both of which will independently be used for alignment summarization in the tax output file. The cov file contains a complete set of total, unique, and Adamantine breadth-of-coverage and depth-of-coverage (i.e., pileup) statistics for each genome or reference sequence, separated into total, unique, and Adamantine k-mer coverage. Because alignment coordinates are not stored, these values are approximated via k-mer set coverage and multiplicity of coverage per reference for each k-mer type (e.g., total Adamantine coverage, total unique coverage, total coverage).

Depth of coverage can be interpreted as read pileup, which can be used to generate relative abundance estimates. The ref file contains a simple table of references matched, and the number of queries assigned to each. The perq file will report the potential assignments of every read, enabling users to further understand their alignment results (e.g., filtering multi-mapping reads). Finally, the orthog file (--orthog-out) cross-tabulates the winning assignments from the primary and secondary taxonomic hierarchies. Note that each output type is only computed if it is requested by its corresponding output flag (e.g., --cov-out), to avoid unnecessary bookkeeping.

By default, XTree does not threshold based on coverage in order to determine genome presence and absence. While this step is critical for determining meaningful compositional analysis downstream, the decision is left up to the user, as the selected threshold will vary based on the sample set and tolerance to noise. Based on the analysis in this manuscript, we recommend a 1% threshold derived from the minimum of total, unique, and Adamantine coverage. Recall that, for example, 1% Adamantine coverage does not correspond to covering 1% of the genome; it instead indicates 1% coverage of the ultra-unique (Adamantine) regions of the genome. We provide scripts for assisting with coverage thresholding in the GitHub repository.

### XTree pseudocode

Below is pseudocode for the XTree algorithm:

Inputs:

1. Reference genomes G = {g_1_ … g_N_}
2. Query reads Q = {q_1_ … q_M_}
3. k-mer length K (BUILD mode)
4. Prefix length PL (BUILD mode)
5. Compression level C (BUILD mode)
6. Optional taxonomy mapping T (BUILD mode) <u>BUILD:</u>

Construct lookup data structure PrefixBins with length = all possible prefixes

1. For each reference genome g_i_ in G:

a. Stream through g_i_ and extract canonical k-mers, defined as the lexicographically smaller of the forward k-mer and its reverse complement.
b. Apply motif-based (C ≤ 2) or hash-based (C ≥ 3) subsampling according to compression level C.
c. For each retained k-mer, split it into a prefix pfx of length PL and a suffix sfx.
d. Append the pair (sfx, i) to the bin PrefixBins[pfx][] corresponding to prefix pfx.

For each prefix bin:

- Sort all stored (sfx, i) pairs lexicographically by sfx, then by reference index i.

If Adamantine is enabled, construct the Adamantine disqualifier tree:

1. Initialize a new prefix-bin data structure WeakBins with length = all possible prefixes.
2. For each reference genome g_i_ in G:

a. Identify unique k-mers by examining multiplicity of suffixes within each prefix bin in PrefixBins.
b. Mark an indexed, reference-unique k-mer as weak if a k-mer from any other reference differs from it by one substitution on either strand. The remaining reference-unique k-mers are Adamantine.

Store the database as:

- A prefix offset array mapping each prefix to a contiguous suffix range.
- A flat, suffix-sorted array of (suffix, reference index) pairs.
- Reference metadata and optional taxonomy mappings.
- Optionally, the Adamantine disqualifier tree, by flattening WeakBins.

### ALIGN

For each query read q_j_ in Q:

1. Extract canonical k-mers, apply the same subsampling as in BUILD, and split each into (pfx, sfx).

a. For each (pfx, sfx):

i. Locate the suffix range corresponding to pfx.
ii. Left-first binary-search for sfx within that range.
iii. Enumerate all reference indices sharing that suffix.
iv. Aggregate k-mer hit counts per reference for q_j_.
2. If taxonomy output is requested, derive taxonomic labels for q_j_ directly from the set of possible reference hits using lowest-common-ancestor interpolation with a confidence threshold.

a. If shallow LCA selected, use voting scheme among leaves
3. Assign q_j_ to exactly one reference using one of the following modes:

a. BEST: select the reference with the highest k-mer count for q_j_.
b. CAPITALIST: defer assignment for global redistribution.

If coverage output is requested:

1. Tally matched k-mers per reference across all reads.
2. Compute total, unique, and Adamantine coverage statistics.

a. If --doforage is specified, consider all references with at least 1 k-mer match
b. If --half-forage is specified, consider the set of k-mers mapping to the highest supported reference and all others with at least two-thirds of its k-mer support
c. If neither is specified, consider the set of k-mers mapping to the highest supported reference only
3. If CAPITALIST redistribution is enabled:

a. Iteratively reassign reads to references using a global expectation–maximization-like procedure until convergence or the pass limit (seven passes by default) is reached.
b. Output final one-to-one read-to-reference assignments.

Outputs:

- Reference assignments, optional taxonomy summaries, and optional coverage statistics.

### Construction of XTree prokaryotic reference databases

For the prokaryotic benchmark, we constructed XTree databases from the representative genomes of 20,000 species clusters randomly selected from the Genome Taxonomy Database (GTDB) Release 232 (see below); 19,999 of these genomes were present in the final reference set and database, because one genome failed to download from NCBI. All benchmarks used XTree v2.2.13. For the k-mer sweep (Supp Fig 1), we additionally indexed these genomes with XTree’s BUILD function at k = 21 and 24 at compression levels 2, 4, 8, and 12, and aligned and scored both read sets exactly as described below for k = 29 (whose results at the same compression levels were taken from the main benchmark). Pre-indexed databases for GTDB Releases 214–226 are provided separately.

### Construction of synthetic prokaryotic metagenomes

To evaluate the performance of XTree, we constructed synthetic metagenomes. We randomly selected 20,000 GTDB Release 232 species clusters that contained at least one representative and one non-representative genome; because clusters were drawn from all of GTDB, 18,988 were bacterial and 1,012 archaeal. A set containing the 20,000 representative genomes was then created, and a second set of 20,000 corresponding non-representative genomes was made by randomly selecting one of the non-representative genomes within each representative genome cluster.

We then generated 100 synthetic metagenomes each comprising 75 to 100 prokaryotic genomes from the previously generated genome sets. The first 50 metagenomes (training set) were created using randomly selected genomes from the representative genomes. The final 50 metagenomes (testing set) were made using non-representative genomes. For each of the metagenomes, every genome was then assigned an abundance (i.e., depth of coverage) ranging from low abundance (0.01X coverage) to high abundance (5X coverage). The abundances were randomly assigned following a log-uniform distribution with many low abundance genomes and few high abundance genomes.

### Short-read and Long-read Simulation

To generate synthetic Illumina sequencing reads for each metagenome, we used DWGSIM^24^ with a per-base error rate of 0.001 and 150-bp paired-end reads. Each synthetic metagenome was created by concatenating reads from all genomes in a given set.

Next, a similar approach was taken with long-read data. Synthetic ONT-HQ reads for each metagenome were simulated using PBSIM3^25^. The read lengths varied from 500–10,000 bp (mean 4,000 bp, SD 3,000 bp) with an accuracy of 99.5%, again with assigned abundances following a log-uniform distribution from 0.01X to 5X coverage. For each synthetic metagenome, the reads were again concatenated from all genomes within the given set.

### XTree Alignment and Compression Benchmarking

To test the performance of XTree (v2.2.13), the BUILD function was run on the representative genomes with k = 29 and 16 threads (--threads 16) at each compression level from 15 down to 1 (compression level 0 was not evaluated). After each build, the simulated Illumina reads for each of the 50 training and 50 testing metagenomes were aligned using XTree ALIGN with default settings (i.e., without Capitalist redistribution) except --threads 16, writing coverage (--cov-out), per-query (--perq-out), and per-reference (--ref-out) outputs. The same process was then repeated for the ONT-HQ long-read metagenomes against the same database. For each read type and compression level, one untimed warm-up alignment preceded the timed alignments to load the database into the page cache.

For this and all other synthetic metagenome evaluation tests, expected relative abundance was equal to the simulated coverage (DWGSIM or PBSIM3) for a given genome, normalized to sum to one. Observed relative abundance was the similarly normalized depth of coverage (pileup; the Coverage_est column of the cov output) reported by XTree; for testing metagenomes, expected abundances were summed by species-cluster representative. Agreement was summarized as the Pearson correlation between expected and observed relative abundance across all genomes in each set. A genome was called present when the minimum of its total, unique, and Adamantine proportions covered (min(Proportion_covered, Unique_proportion_covered, Adamantium_prop) in the cov output) exceeded a coverage threshold. We report results at a fixed 1% threshold, and additionally swept thresholds from 0 to 10% in 0.1% increments to identify the F1-maximizing threshold at each taxonomic rank and compression level. For testing metagenomes, each non-representative genome was mapped to its species-cluster representative using the GTDB Release 232 cluster file, so that detection of that representative counted as a true positive. At ranks above species, calls were aggregated to the GTDB lineage at that rank, with each taxon assigned the maximum coverage of its member genomes. Precision, recall, and F1 scores were calculated per metagenome and averaged over the 50 metagenomes in each set at each of the 15 compression levels; ± values denote SD unless otherwise stated.

For the archaeal benchmark, we repeated the same procedure using the 3,013 GTDB Release 232 archaeal species clusters that contained a representative and at least one non-representative genome (3,012 representative genomes were present in the final database, because one genome failed to download from NCBI). Synthetic short- and long-read metagenomes (50 training and 50 testing) were generated with the same scripts and parameters as for the prokaryotic benchmark, the database was built with XTree v2.2.13 at k = 29 at each compression level from 15 down to 1, and alignment and scoring followed the prokaryotic benchmark exactly, using the archaeal subset of the GTDB Release 232 cluster file to map non-representative genomes to their representatives.

### Per-order coverage thresholds

To assess how F1-optimal coverage thresholds vary across clades (Fig 3), we used the compression level 4 database (XTree v2.2.13, k = 29) and the short and long-read metagenomes of both sets. Each reference genome was assigned to its GTDB Release 232 order, and the truth positives (with non-representative genomes mapped to their cluster representatives, as above) and the references reported by XTree were grouped by order. For each order, we swept thresholds from 0 to 100% in 0.1% increments separately on total coverage (Proportion_covered), unique coverage (Unique_proportion_covered), and Adamantine coverage (Adamantium_prop), calling a reference present when its coverage exceeded the threshold. Because most orders contribute only a few genomes to any one metagenome, true positives, false positives, and false negatives were pooled across the 50 metagenomes of each set before computing F1, and the F1-maximizing threshold was taken as the lowest threshold achieving the maximum F1. To compare the fixed 1% threshold with each order’s optimum, we also recorded F1 and the number of false positives at 1%. We report orders with at least 20 positive genome–metagenome pairs in a set (42 orders in the testing set and 40 in the training set).

### Benchmarking of XTree against Other Aligners

To compare the performance of XTree against other tools commonly used in metagenomic profiling workflows, we ran Kraken2 with Bracken, Ganon2, Centrifuge, KMCP, and MetaPhlAn on the same sets of synthetic metagenomes. Kraken2, Ganon2, Centrifuge, and KMCP were run with default settings, each building a database from the same representative genomes used for XTree and then classifying the 100 short-read and 100 long-read synthetic metagenomes. MetaPhlAn was run with its default marker-gene database, using Bowtie2 to align short reads and Minimap2 to align long reads. Taxa reported by each tool under its default presence call were scored at the species level against the same ground truth as XTree. For KMCP, runtime was the sum of the search and profile steps. For XTree, we used the compression level 4 database (the v2.2.13 default) with the 1% coverage threshold.

### Evaluation of runtime and memory usage

In order to determine the effects of compression level on XTree’s performance and how it performed relative to other aligners, we tracked runtime and memory usage across both database-building and alignment/classification stages for each XTree compression level and each tool. Wall-clock time was recorded for each run, and peak memory was recorded by polling each process’s resident set size (VmRSS and VmHWM in /proc) every 0.05 s. Because XTree memory-maps its database, its peak resident set size during alignment includes the shared, file-backed database pages. All testing was performed on Google Cloud Compute Engine virtual machines of a single machine type, n4d-custom-16-262144-ext (N4D family; 16 vCPUs and 256 GB of RAM), running Debian GNU/Linux 13; every XTree run (including the k-mer sweep and the archaeal benchmark) and every comparison tool used this configuration with 16 threads. A Hyperdisk Balanced volume (10,000 IOPS; 1,000 MB/s throughput) was attached. For Centrifuge, the machine was re-instanced as n4d-custom-16-524288-ext (identical except for memory extended to 512 GB) because its database build exceeded 256 GB (peak resident set size 385 GB) and failed with an out-of-memory error on the 256 GB machine; all other tools ran within 256 GB. For the tool comparison (Fig 2B–C), the first two metagenomes of each set were excluded from runtime averages because they load the database into memory from disk; mean runtime per metagenome was computed over the remaining 96 runs, and error bars show the standard error of that mean. Peak alignment memory is the maximum over all 100 runs.

## Data Availability

All code used in this study can be accessed through the Two Frontiers Project XTree GitHub repository (https://github.com/two-frontiers-project/2FP-XTree).

## Acknowledgements

We thank Heng Li and Jim Shaw for their feedback on an earlier version of this manuscript.

---

**Supplemental Figure 1:**
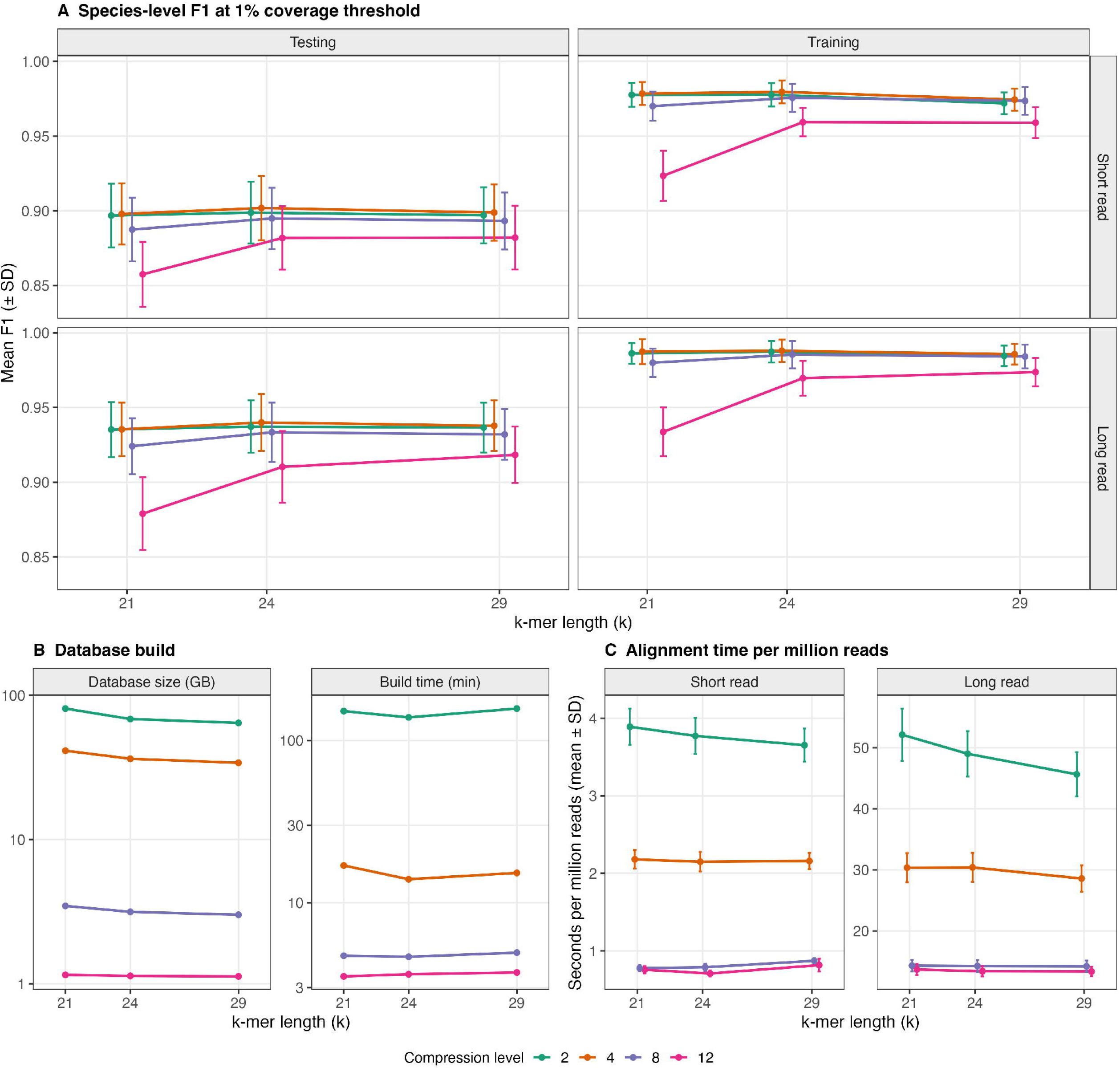
Effect of k-mer length on XTree performance (XTree v2.2.13). Databases were built at k=21,24 and 29 at compression levels 2, 4, 8 and 12 and evaluated, with k=29 values being used from the main benchmark. A) Species-level mean F1 score at a 1% coverage threshold for all synthetic metagenomes (training and testing, short and long reads). B) Database size and build time (Log scale). C) Mean alignment (± SD; testing and training runs combined) time per 1 million reads.

**Supplemental Figure 2:**
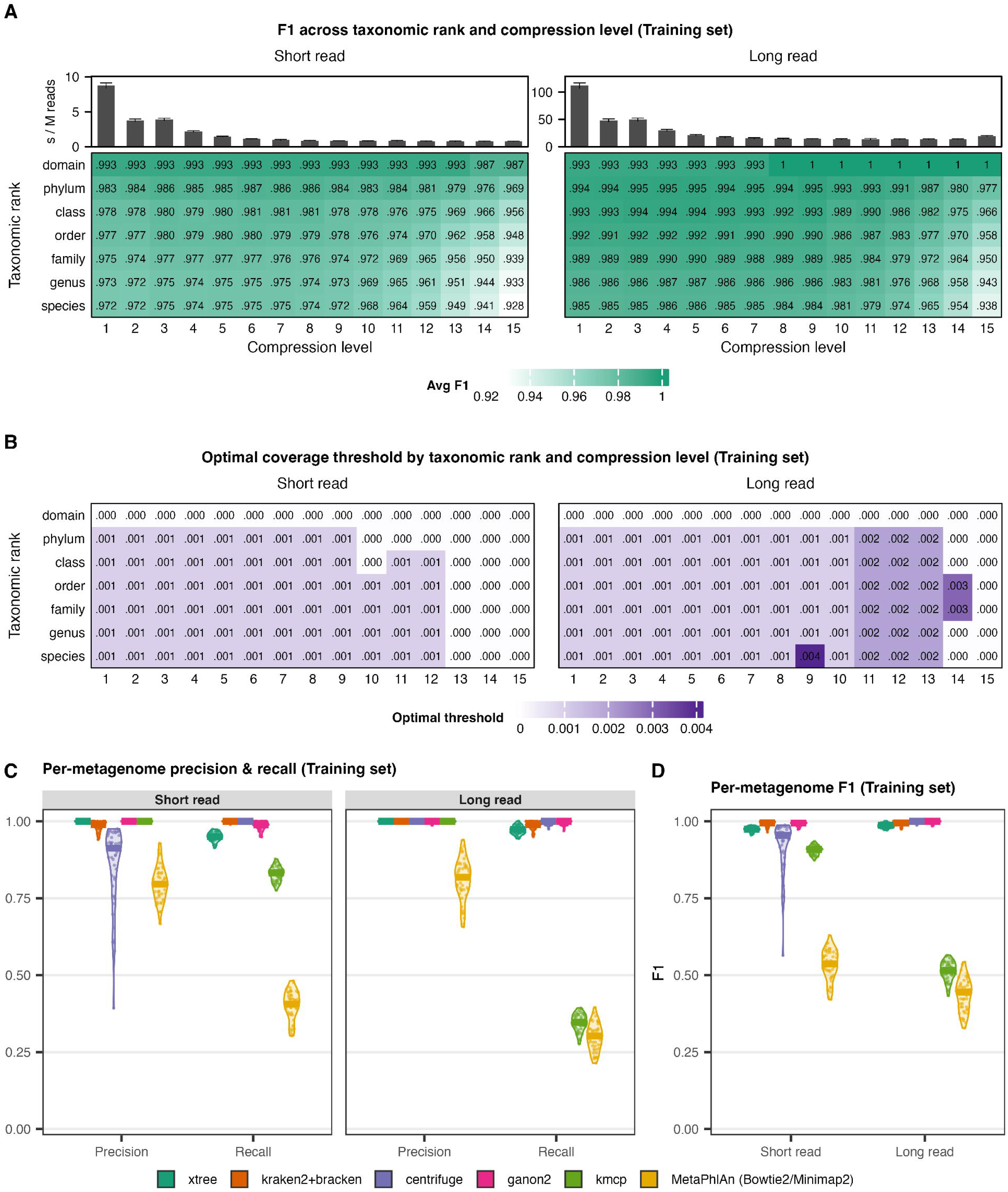
XTree performance on the training set of synthetic metagenomes. The training set comprises the 50 synthetic metagenomes built from reference genomes indexed during database formation. A) Mean F1 score by taxonomic rank and compression level at a 1% coverage threshold, for short and long reads, with mean alignment time per million reads. B) Coverage threshold that maximizes mean F1 score at each taxonomic rank and compression level. C) Per-metagenome precision and recall of XTree and other profilers. D) Per-metagenome species-level F1 score. In panel C and D, each point is one metagenome and horizontal lines mark the median. XTree was used at compression level 4 (k=29) and a 1% coverage threshold. The other tools were run with default settings, and MetaPhlAn4 using Bowtie2 for short reads and Minimap2 for long reads.

**Supplemental Figure 3:**
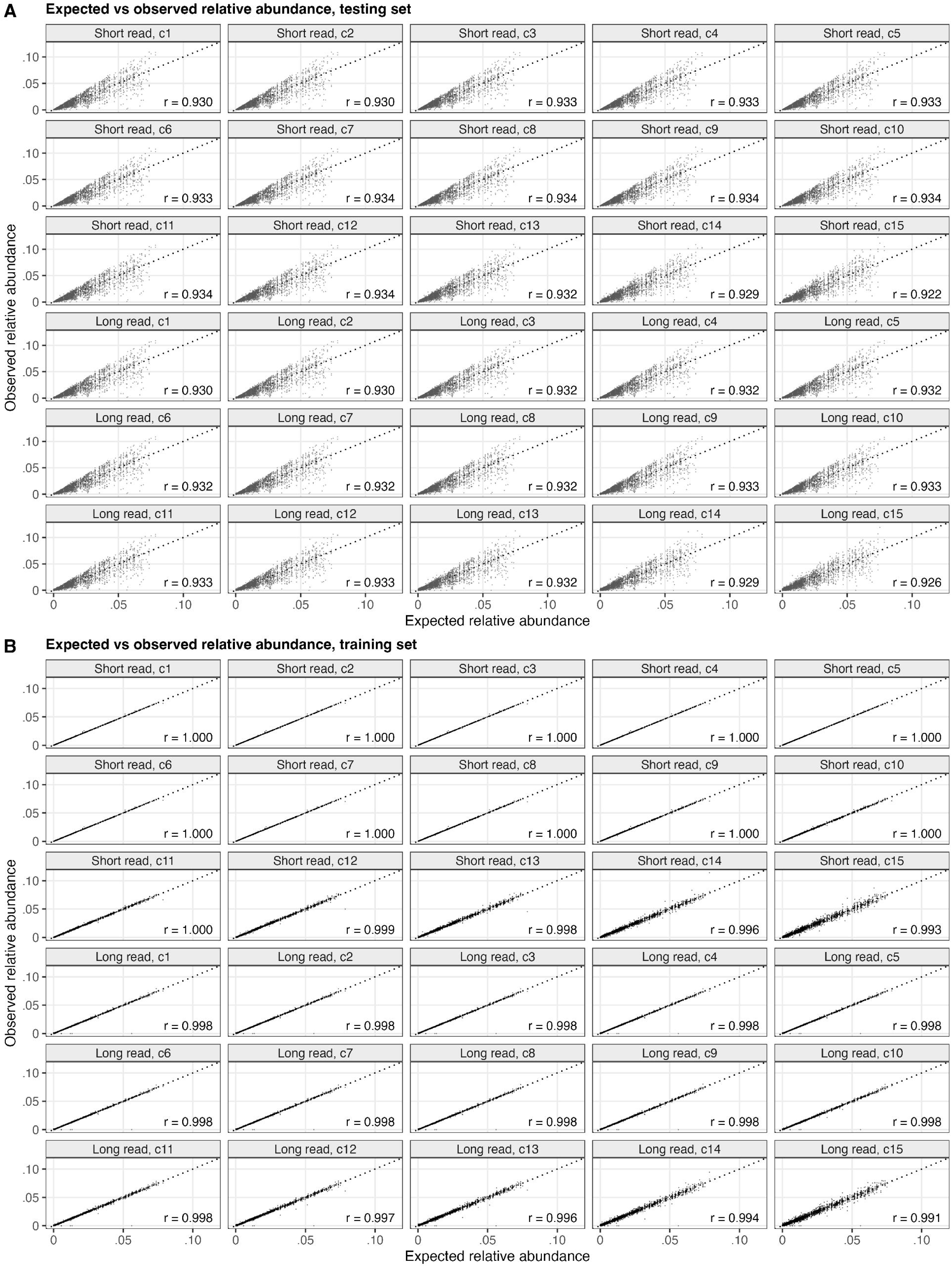
Expected versus observed relative abundance at each compression level. A) Testing set. B) Training set. Each panel shows one read type and compression level (C1–C15), rows 1-3 are short reads and rows 4-6 are long reads. Each point is one genome in one metagenome. Expected relative abundance comes from the simulated coverage (truth data). Observed relative abundance is XTree’s estimated coverage depth for that genome divided by the total over all reported genomes for that metagenome. The dotted line marks y=x, the perfect scenario, and each panel gives the Pearson correlation (r).

**Supplemental Figure 4:**
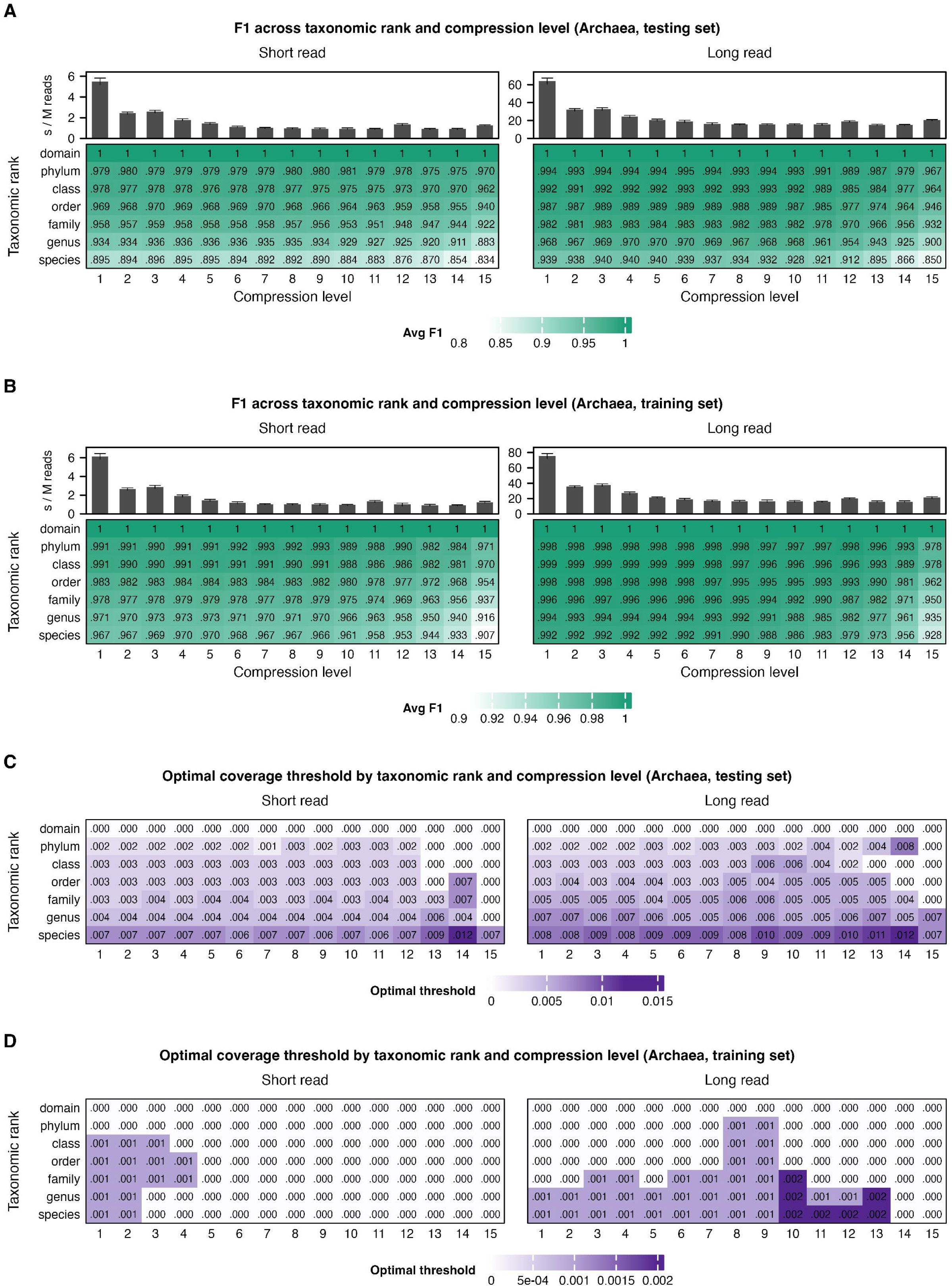
XTree performance on synthetic archaeal metagenomes. Databases were built with XTree v2.2.13 at k=29 from 3,012 GTDB archaeal representative genomes. A-B) Mean F1 score by taxonomic rank and compression level at a 1% coverage threshold for the 50 testing (A) and 50 training (B) metagenomes for short and long reads, with bars above each heatmap showing mean alignment time per million reads. C-D) Coverage threshold that maximizes mean F1 score at each taxonomic rank and compression level for the testing (C) and training (D) sets.

**Supplemental Figure 5:**
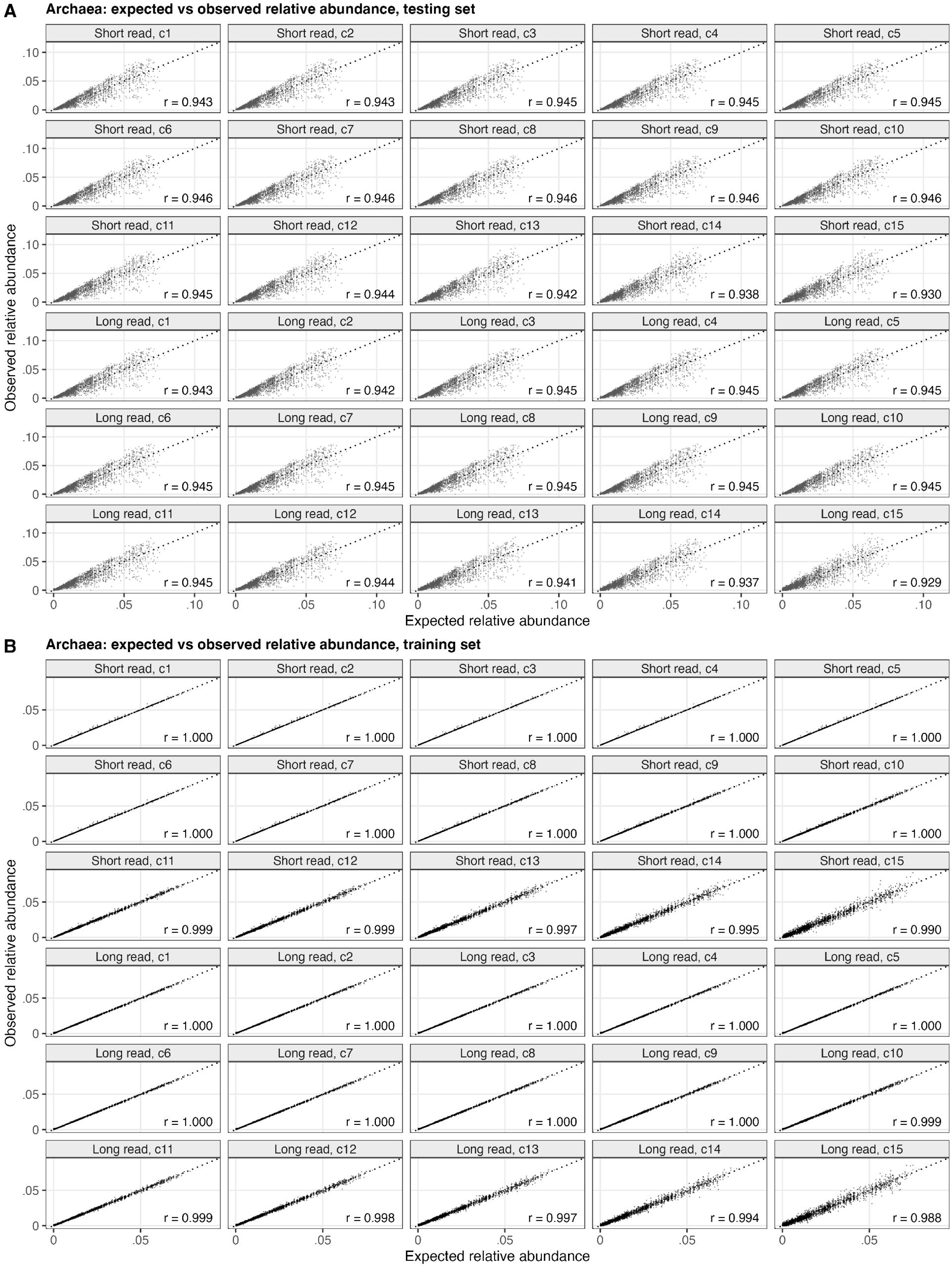
Expected versus observed relative abundance for synthetic archaeal metagenomes. A) Testing set. B) Training set. The layout and definitions are as in Supplemental Figure 3.

## References

1. Al-Ghalith, G. & Knights, D. BURST enables mathematically optimal short-read alignment for big data. Preprint at 10.1101/2020.09.08.287128 (2020).

2. Priya, S. et al. Identification of shared and disease-specific host gene–microbiome associations across human diseases using multi-omic integration. Nat. Microbiol. 7, 780–795 (2022).

3. Tierney, B. T. et al. Longitudinal multi-omics analysis of host microbiome architecture and immune responses during short-term spaceflight. Nat. Microbiol. 9, 1661–1675 (2024).

4. Button, J. E. et al. Precision modulation of dysbiotic adult microbiomes with a human-milk-derived synbiotic reshapes gut microbial composition and metabolites. Cell Host Microbe 31, 1523–1538.e10 (2023).

5. Tolonen, A. C. et al. Synthetic glycans control gut microbiome structure and mitigate colitis in mice. Nat. Commun. 13, 1244 (2022).

6. Lin, C.-Y. et al. Longitudinal fecal microbiome and metabolite data demonstrate rapid shifts and subsequent stabilization after an abrupt dietary change in healthy adult dogs. *Anim*. Microbiome 4, 46 (2022).

7. Johnson, K. E. et al. Human cytomegalovirus in breast milk is associated with milk composition and the infant gut microbiome and growth. Nat. Commun. 15, 6216 (2024).

8. Vinithakumari, A. A. et al. Clostridioides difficile infection dysregulates brain dopamine metabolism. Microbiol. Spectr. 10, e0007322 (2022).

9. Tierney, B. T. et al. Towards geospatially-resolved public-health surveillance via wastewater sequencing. Nat. Commun. 15, 8386 (2024).

10. Napier, B. A. et al. Mo1894 A multi-species synbiotic (DS-01) alleviates constipation and abdominal pain in irritable bowel syndrome subtype mixed (IBS-M) subjects while boosting synbiotic species associated with decreased systemic inflammation and NET formation. Gastroenterology 166, S-1164 (2024).

11. Wood, D. E., Lu, J. & Langmead, B. Improved metagenomic analysis with Kraken 2. Genome Biol. 20, 257 (2019).

12. Irber, L. et al. sourmash v4: a multitool to quickly search, compare, and analyze genomic and metagenomic data sets. J. Open Source Softw. 9, 6830 (2024).

13. Roberts, M., Hayes, W., Hunt, B. R., Mount, S. M. & Yorke, J. A. Reducing storage requirements for biological sequence comparison. Bioinformatics 20, 3363–3369 (2004).

14. Parks, D. H. et al. GTDB release 10: a complete and systematic taxonomy for 715 230 bacterial and 17 245 archaeal genomes. Nucleic Acids Res. 10.1093/nar/gkaf1040 (2025).

15. Lu, J., Breitwieser, F. P., Thielen, P. & Salzberg, S. L. Bracken: estimating species abundance in metagenomics data. PeerJ Comput. Sci. 3, e104 (2017).

16. Langmead, B. & Salzberg, S. L. Fast gapped-read alignment with Bowtie 2. Nat. Methods 9, 357–359 (2012).

17. Piro, V. C. & Reinert, K. ganon2: up-to-date and scalable metagenomics analysis. NAR Genom. Bioinform. 7, lqaf094 (2025).

18. Shen, W. et al. KMCP: accurate metagenomic profiling of both prokaryotic and viral populations by pseudo-mapping. Bioinformatics 39, btac845 (2023).

19. Kim, D., Song, L., Breitwieser, F. P. & Salzberg, S. L. Centrifuge: rapid and sensitive classification of metagenomic sequences. Genome Res. 26, 1721–1729 (2016).

20. Blanco-Míguez, A. et al. Extending and improving metagenomic taxonomic profiling with uncharacterized species using MetaPhlAn 4. Nat. Biotechnol. 41, 1633–1644 (2023).

21. Li, H. Minimap2: pairwise alignment for nucleotide sequences. Bioinformatics 34, 3094–3100 (2018).

22. Steele, G. L., Lea, D. & Flood, C. H. Fast splittable pseudorandom number generators. In Proc. 2014 ACM International Conference on Object Oriented Programming Systems Languages & Applications (OOPSLA ’14) 453–472 (ACM, 2014).

23. Farach-Colton, M. Lowest common ancestors in trees. In Encyclopedia of Algorithms (ed. Kao, M.-Y.) 1169–1174 (Springer, 2016).

24. Homer, N. DWGSIM: whole genome simulator for next-generation sequencing. GitHub https://github.com/nh13/DWGSIM.

25. Ono, Y., Hamada, M. & Asai, K. PBSIM3: a simulator for all types of PacBio and ONT long reads. NAR Genom. Bioinform. 4, lqac092 (2022).

